# Activation mechanism of the cardiac calcium pump by a small-molecule allosteric modulator

**DOI:** 10.1101/2023.09.07.556734

**Authors:** Jaroslava Šeflová, Carlos Cruz-Cortés, Guadalupe Guerrero-Serna, Seth L. Robia, L. Michel Espinoza-Fonseca

**Affiliations:** Center for Arrhythmia Research, Department of Internal Medicine, Division of Cardiovascular Medicine, University of Michigan, Ann Arbor, MI 48109, USA; Department of Cell and Molecular Physiology, Loyola University Chicago, Maywood, IL 60153, USA

## Abstract

The discovery of small-molecule allosteric modulators is an emerging paradigm in drug discovery, and signal transduction is a subtle and dynamic process that is challenging to characterize. We developed a time-correlated single photon counting (TCSPC) imaging approach to investigate the activation mechanism of a druggable protein by a small-molecule allosteric modulator. We tested this approach using the cardiac sarcoplasmic reticulum Ca^2+^-ATPase (SERCA2a), an important pharmacological target that transports Ca^2+^ at the expense of ATP hydrolysis in the heart. We found that CDN1163, a validated SERCA2a activator, does not dissociate the endogenous complex between SERCA2a and its regulator phospholamban (PLN) in the presence of either Ca^2+^ or AMP-PCP, a non-hydrolyzable ATP analog. CDN1163 does not influence SERCA2a’s affinity for Ca^2+^ ions at functionally relevant conditions. Global analysis of the fluorescence lifetimes showed that ATP is both a substrate and a modulator that populates competent SERCA2a conformations. Interestingly, CDN1163 alone does not significantly induce changes in the structural populations of SERCA2a. Instead, CDN1163 potentiates the effects of ATP to further shift the equilibrium toward a competent SERCA2a conformation. Importantly, this population shift occurs at sub-physiological conditions, and within physiological Ca^2+^ concentrations at which SERCA2a operates. We propose an activation mechanism whereby a small-molecule modulator synergizes with ATP to stabilize a conformation of SERCA2a primed for activation. This study demonstrates the power of TCSPC to reveal novel insights into how structural and biochemical states are coupled to allosterically activate a pharmacological target in the heart.

## Introduction

The concept of allostery for drug discovery is an evolving paradigm in medicinal chemistry.^1-2^ In this paradigm, allosteric drugs modulate the activity through the propagation of allosteric signaling, producing either inhibition or activation of the target. Understanding the structural mechanisms for small-molecule modulation is an essential prerequisite for allosteric drug discovery,^3^ but signal transduction is a dynamic process involving subtle structural changes^4-5^ that are challenging to characterize using traditional experimental techniques.^6^ This motivates the development of novel approaches to determine the structural mechanisms underlying small-molecule allosteric modulation of druggable proteins.

The cardiac calcium pump (Sarcoplasmic reticulum Ca^2+^-ATPase, SERCA2a) plays an essential role in normal cardiac function, clearing cytosolic Ca^2+^ needed to relax muscle cells in each heartbeat (diastole).^7^ SERCA pumps two Ca^2+^ ions into the SR lumen using energy derived from the hydrolysis of ATP^8-9^ and is regulated by the 52-residue membrane protein phospholamban (PLN). PLN regulates SERCA2a by inhibiting its ability to transport Ca^2+^,^10-11^ and inhibition is relieved by phosphorylation of PLN.^12-16^ A key molecular dysfunction in patients with heart failure usually involves insufficient SERCA expression and impaired PLN phosphorylation, leading to SERCA2a inactivation and decreased Ca^2+^ transport in the cardiomyocyte. Reactivation of Ca^2+^ transport results in improved cardiac function in experimental models of heart failure,^17-21^ and SERCA2a is a well-validated target and its small-molecule activation is a promising approach for heart failure therapy.^22-23^

A conventional concept in the field is that dissociation of the endogenous regulatory SERCA2a–PLN complex is a requirement for activation of SERCA2a,^24-25^ and small molecules that compete with PLN have been reported in the literature.^26-28^ This concept has been recently challenged by spectroscopy experiments showing that relief of SERCA2a–PLB inhibition occurs primarily by structural changes within the regulatory complex, not by dissociation of PLN.^29^ Despite the breakthrough discovery of novel small-molecule modulators of SERCA2a ^30-31^ and the advances in the structural biology of SERCA,^32^ there are no crystal/cryo-EM structures of SERCA2a to allosteric activators, hence the molecular basis for smallmolecule activation of the pump remains unknown. In this study, we developed a time-correlated single photon counting (TCSPC) imaging method to investigate in unprecedented detail the mechanisms for SERCA2a activation by a small-molecule allosteric modulator. The result is a vivid visualization of the molecular mechanism for the allosteric modulation of a druggable target in the heart.

### Experimental section

#### Chemicals

All chemicals used in this study were purchased at reagent quality (purity>95% by HPLC): CDN1163, *N*-(2-methylquinolin-8-yl)-4-propan-2-yloxybenzamide (Sigma, St. Louis, MO); istaroxime, (3*E*,5*S*,8*R*,9*S*,10*R*,13*S*,14*S*)-3-(2-aminoethoxyimino)-10,13-dimethyl-1,2,4,5,7,8,9,11,12,14,15,16-dodecahydrocyclopenta[a]phenanthrene-6,17-dione (Med-ChemExpress LLC, Monmouth Junction, NJ); CP-154526, *N*-butyl-*N*-ethyl-2,5-dimethyl-7-(2,4,6-trimethylphenyl)pyrrolo[2,3-d]pyrimidin-4-amine (Sigma, St. Louis, MO), Ro 41-0960, (3,4-dihydroxy-5-nitrophenyl)-(2-fluorophenyl)methanone (Sigma, St. Louis, MO); AMP-PCP, β,γ-Methyleneadenosine-5’-triphosphate (Sigma, St. Louis, MO).

#### Isolation of enriched SERCA2a microsomes

Pig hearts were obtained after euthanasia and placed in a cardioplegic solution (280 mM glucose, 13.44 mM KCl, 12.6 mM NaHCO_3_, and 34 mM mannitol). Left ventricles free walls were obtained, minced, and homogenized with a cold buffer that contained 9.1 mM NaHCO_3_, 0.9 mM Na_2_CO_3,_ and a cocktail of proteases inhibitors (Sigma, St. Louis, MO); the mixture was centrifuged at 6,500 *g* for 30 minutes at 4°C to remove debris. The supernatant was filtered, collected, and centrifuged at 14,000 *g* for 30 min at 4°C. The collected filtrate was centrifuged at 47,000 *g* for 60 min at 4°C. The pellet was resuspended in a solution containing 0.6 M KCl and 20 mM Tris (pH=6.8). The suspension was centrifuged at 120,000 *g* for 60 min at 4°C, and the pellet was resuspended in a solution containing 0.3 M sucrose, 5 mM MOPS, and protease inhibitors (pH=7.4). The protein concentration of the SR microsomal fraction was determined using the PierceTM Coomassie plus assay kit (Thermo-Fisher Scientific, Waltham, MA). The microsomal membranes were aliquoted, frozen in liquid nitrogen, and stored at -80°C.

#### SERCA ATPase activity assays

We performed SERCA2a activity assays using an enzyme-coupled NADH-linked ATPase assay described previously.^33^ Briefly, we measured the activity of Ca^2+^ ATPase in µmol min^-1^ mg^-1^ from the decrease in absorbance of NADH at 340 nm at 25°C in a 96-well format using a Synergy H1 (BioTek, Winooski, VT) microplate reader. Each well contained a 200 µl final volume of assay buffer containing SERCA buffer (50 mM MOPS, 100 mM KCl, 5 mM MgCl_2_, and 1 mM EGTA, pH=7), 5U lactate dehydrogenase, 5U pyruvate dehydrogenase, 1 mM phosphoenolpyruvate, 5 mM ATP, 0.2 mM NADH, 2 µg of microsomal suspension, 2 µM of Ca^2+^ ionophore A23187, and eight free Ca^2+^ concentrations. Each concentration of the compounds tested here was calculated to a final volume of 200 µl. The small molecules were incubated for 30 min at 25°C with the reaction mixture. Concentration-response curves for each compound were constructed with the data from [Ca^2+^]-dependent SERCA activity curves performed at compound concentrations of 0.1-100 µM. Each plate included untreated and thapsigargin-treated microsomes as controls. In all cases, the maximal activity values (V_max_) were normalized relative to the compound-free samples.

#### HEK-293T cells culture

The human two-color SERCA2a construct was produced as described previously.^34^ The mCerulean at the N-terminus of human SERCA2a was replaced by mCyRFP1^35^ and YFP was replaced by mMaroon1^36^ at the position before residue number 509 within the N-domain of SERCA2a (**Fig. 1A**). For microsomes containing SERCA2a and PLN, the mCyRFP1 and mMaroon were fused to the N-termini of SERCA2a and PLN,^37^ respectively. The mCyRFP1-mMaroon1 construct has a Förster distance of 63.34 Å.^35, 38^ The final constructs were verified by full-length sequencing (ACGT, USA). HEK 293T cells (CRL-3216, ATCC) were seeded into a 150 mm cell culture dish 72 hours before transient transfection and cultured in DMEM (Corning, USA) supplemented with 10% FBS in 5% CO_2_ incubator at 37°C. When cells reached 70 – 80% confluence, the cell culture media was replaced by fresh media, and cells were transfected with two-color SERCA construct^34, 38^ using Lipofectamine 3000 kit (Invitrogen, Life Technologies). Cells were harvested 48 hours post-transfection and microsomal membranes were prepared on the day of the harvest.

**Fig. 1.**
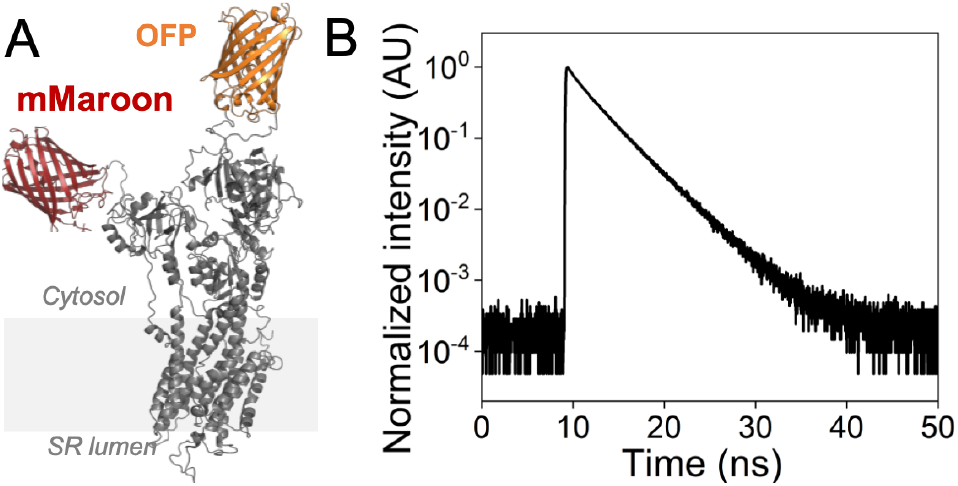
(A) Structural representation of the two-color cardiac SERCA2a construct labeled with maroon fluorescent protein (mMaroon) at position Met1 and with orange fluorescent protein (OFP) at position Gly509. The approximate location of the lipid bilayer represented by the shaded gray box. (B) Fluorescence decay measured by TCSPC for two-color SERCA2a in DMSO vehicle. We found that the TCSPC imaging approach has over 4 orders of magnitude difference between signal (intensity at ∼10^0^ AU) and noise (intensity at ∼10^-4^ AU).

#### Microsomal membranes preparation from transfected HEK-293T cells

The microsomal membranes from HEK-293T were prepared according to the protocol published previously.^34, 39^ Specifically, each 150 mm cell culture dish was scraped into 12 ml of homogenization buffer (0.5 mM MgCl2, 10 mM Tris-HCl; pH 7.5) supplemented with UltraCruz Protease Inhibitor Cocktail Tablet without EDTA (Santa Cruz Biotechnology). Cells were pelleted using centrifugation of 1,000 *g* for 10 minutes at 4°C and subsequently dissolved in 5 ml of fresh homogenization buffer. The cell suspension was homogenized with 20 strokes of Potter-Elvehjem glass homogenizer, and 5 ml of sucrose buffer (100 mM MOPS, 500 mM sucrose; pH=7.0 with protease inhibitors). The crude homogenate was passed 10 times through a 27-gauge needle. Subsequently, the homogenate was centrifuged at 1,000 *g* for 10 minutes at 4°C to remove unbroken cells, mitochondria, and cellular debris. The supernatant was subjected to ultracentrifugation at 126,000 *g* for 30 minutes at 4°C, and the pellet was dissolved in a 1:1 mixture of homogenization and sucrose buffers. The total protein concentration was determined using BCA assay (Pierce BCA Protein assay, ThermoFisher Scientific, Rockford, IL). The microsomal fractions were separated into single-use aliquots containing 50 μl of microsomal membranes at a concentration of 3 – 4.5 mg/ml. We performed four independent transfections to account for biological variability as part of our experimental design.

#### PLN-CDN1163 competition experiments

The SERCA2a competition experiment monitored FRET changes between mCyRFP1-SERCA2a and mMaroon1-PLN microsomes. We used a physiologically relevant SERCA-to-PLN DNA ratio of 1:5,^40^ and tested the effects of CDN1163 at increasing concentrations of the compound (0.1–50 μM). The donor alone lifetime was shortened to 2.63 ns due to FRET. This corresponds to average FRET efficiency of 26% using the following equation 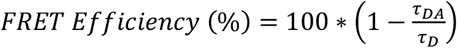, where τ_D_ is the lifetime of donor alone and τ_DA_ is the lifetime of the donor in the presence of acceptor.

#### Time-correlated single-photon counting imaging of SERCA2a

Aliquots of microsomes were thawed on ice, mixed with the vehicle (10 μM DMSO) or 10 µM CDN1163 (>98% purity by HPLC; Sigma-Aldrich), and incubated for 30 minutes at room temperature. Subsequently, the microsomes were mixed with a buffer composed of 120 mM potassium aspartate, 15 mM KCl, 5 mM KH_2_PO_4_, 0.75 mM MgCl_2_, 2% dextran, 20 mM HEPES, 2 mM EGTA, and CaCl_2_ 1.7 mM; pH 7.2 with or without 500 μM AMP-PCP and imaged immediately after mixing. The free [Ca^2+^] concentrations were estimated to range from 0.001 to 100 µM.^41^ Time-correlated single-photon counting (TCSPC) experiments were performed as previously described.^37, 42^ Donor (mCyRFP1) fluorescence in the membrane microsomes was excited by a supercontinuum laser beam (Fianium, Southampton, United Kingdom). The donor signal was acquired using an excitation filter 482/18 nm bandpass filter. Fluorescence emission was detected using a 525/50 nm emission bandpass filter. The focused laser was placed inside of microsomes drop, yielding a 100,000 photons/s count rate. Under those conditions, we observed less than 10% photobleaching during the 60 seconds of acquisition time. Fluorescence was detected through a 1.2 N. A. water-immersion objective with a PMA hybrid detector (PicoQuant, Germany) connected to a single photon-counting module (HydraHarp 300, PicoQuant, Germany) with a time channel width of 16 ps. The TCSPC imaging method has over 4 orders of magnitude difference between signal and noise (**Fig. 1B**), making it ideal to capture the subtle structural changes associated with allosteric modulation.

#### Global analysis of the fluorescence lifetimes

Fluorescence decay histograms from 4 different sets of microsomes were analyzed in SymPhoTime 64 software (Picoquant, Germany) with the TCSPC global fitting tool. The donor alone (mCyRFP1-SERCA2a) showed slightly two-exponential decay with relative amplitude of the second component accounting for less than 8%, therefore the donor alone lifetime was estimated as a single exponential decay with a lifetime of 3.56 ns for all tested conditions. The donor in the presence of the acceptor (two-color SERCA2a or mMaroon1-PLN) was best fitted using a two-exponential decay, where the amplitude-weighted average lifetime (τ_avg_) was shorter than the lifetime of the donor alone that is consistent with FRET between donor and acceptor. For the two-color SERCA2a, the τ_avg_ decreased with increasing Ca^2+^ concentrations. Based on our recent work^39^ and to simplify the analysis, we assume the presence of two populations of SERCA2a, open and closed, that is resolved by TCSPC.

#### Statistical analysis

All results are presented as mean ± standard error of the mean (SEM). Significance was evaluated using the Mann–Whitney U test for paired experiments or a two-way analysis of variance (ANOVA) followed by Dunnett’s post-hoc test to analyze differences between the control and multiple treatments. We used 95% confidence intervals around the differences between the groups for the post-hoc test. Two-sided *p* values were used, and α-level <0.05 was considered significant.

## Results

### Validation of CDN1163 as a SERCA2a activator

There are only a handful of small molecules that have been reported to stimulate SERCA activity: CDN1163, an allosteric activator that stimulates the non-muscle SERCA isoform SERCA2b;^43,44^ istaroxime, a molecule that was proposed to stimulate SERCA2a and inhibits the Na^+^/K^+^-ATPase (NKA);^45^ CP-154526, which was reported to increase the maximal activity of muscle SERCA isoforms;^30^ and Ro 41-0960, a small molecule that increases both the activity and Ca^2+^ affinity of the SERCA2a pump^30^ (**Fig. 2A**). The full activation profiles of these molecules have not yet been reported in the literature, therefore we first used ATPase activity assays to establish the concentration-response profiles of these compounds on SERCA2a. In all cases, the SERCA2a activity is reported as the change of V_max_ relative to the activity of SERCA2a alone. When describing the data, we distinguish between ‘observed’ data (i.e., data points) and ‘estimated’ data (i.e., values derived from the fitted model data).

**Fig. 2.**
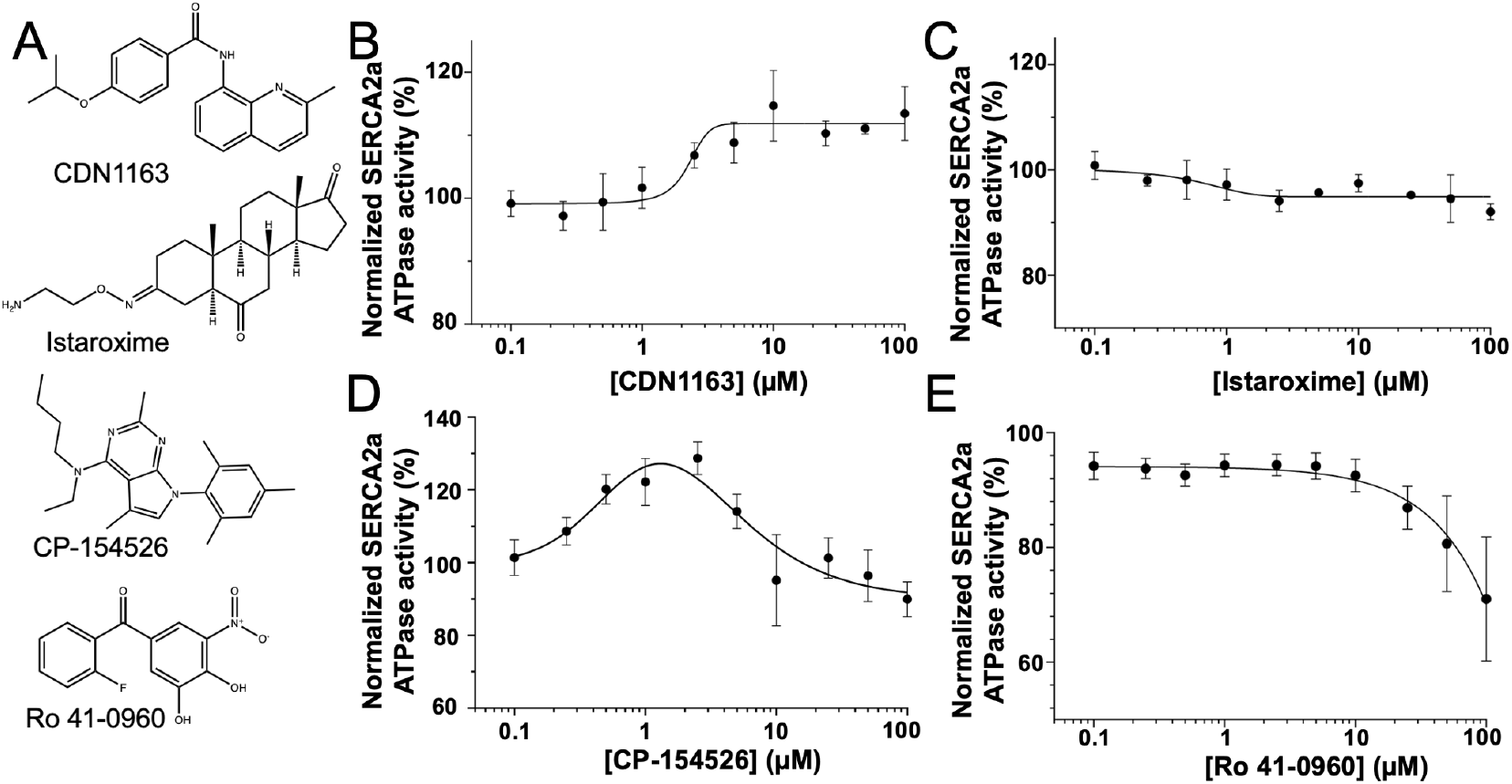
Effects of small-molecule effectors on SERCA2a ATPase activity. (A) Structures of chemically diverse small-molecule effectors reported in the literature as SERCA activators. 10-point concentration-response curves obtained for (B) CDN1163, (C) istaroxime, and (D) CP-154526 and (E) Ro 41-0960 on the ATPase activity of the cardiac SERCA2a. In all cases, the activity of the compounds at each concentration was obtained from an 8-point free Ca^2+^ concentration-dependent SERCA2a activity assay and normalized relative to the untreated control as described in experimental procedures. Data are reported as average ± SEM (N=3).

We found CDN1163 activates SERCA2a in a concentration-dependent manner with an EC_50_ value of 2.3 µM, and a maximal increase in the relative V_max_ activity of 11.8% estimated from the fitted curve (**Fig. 2B**). Notably, we found that the stimulatory effect of CDN1163 on SERCA2a remains constant at compound concentrations >10 µM (**Fig. 2B**). In our hands, istaroxime does not activate SERCA2a at any compound concentration (**Fig. 2C**). At high concentrations, istaroxime has a slight inhibitory effect on SERCA2a; however, this effect is not significant. We speculate that istaroxime does not activate SERCA2a in our assays, probably because of species-specific differences.^26^ However, we note additional ATPase assays performed by us showed that istaroxime inhibits the cardiac isoform of the *α*_1_ isoform of pig Na^+^/K^+^-ATPase (NKA-*α*_1_) with an IC_50_=0.47 µM; this value is virtually identical to that previously reported using the dog NKA-*α*_1_ (IC_50_=0.43 µM).^46^ These findings suggest that, unlike CDN1163, istaroxime may be a species-specific activator of the SERCA2a pump. CP-154526 also activates SERCA2a in the high nM to low µM range, with an observed maximal increase in V_max_ of 28% at a compound concentration of 2.5 µM (**Fig. 2D**). However, this compound follows a bell-shaped dose-response curve, where increasing compound concentration leads to increased activity up to a point and further increases in compound concentration lead to decreasing or abolished activity.^47^ This effect is likely the result of CP154526 aggregation, as suggested before by atomistic molecular dynamics simulations.^48^ The ATPase activation assays showed that Ro 41-0960 does not stimulate the activity of SERCA2a in the concentration range tested here (0.1-100 µM, **Fig. 2E**). Instead, Ro 41-0960 has an inhibitory activity at compound concentrations ≥25 µM, with an observed decrease in V_max_ by ∼30% at a compound concentration of 100 µM (**Fig. 2E**). In summary, we found only CDN1163 activates SERCA2a in a concentration-response manner, hence we used this effector as a molecular probe to study the mechanisms for SERCA2a activation.

### Competitive FRET assays show CDN1163 does not dissociate the SERCA–PLN complex

We used fluorescently labeled SERCA2a and PLN to test whether CDN1163 activates SERCA2a by dissociating the SERCA2a–PLN complex. In this assay, binding of PLN to SERCA2a produces a FRET signal (measured as % of FRET efficiency); dissociation of the complex by CDN1163 is reflected by the decrease of the FRET signal between SERCA2a and PLN (dissociation model)^29^ whereas no changes in the SERCA2a–PLN FRET efficiency suggest that CDN1163 binds to the regulatory complex (subunit model)^29^ (**Fig. 3A**). Activation of SERCA2a requires both Ca^2+^ and ATP. Therefore, we performed these studies at low (0.01 µM) and high (10 µM) free Ca^2+^ concentrations, as well as in the presence and absence of the non-hydrolyzable ATP analog AMP-PCP to account for the effects of Ca^2+^ and ATP. At low Ca^2+^, there is no change in the FRET efficiency between SERCA2a and PLN in the presence or absence of AMP-PCP at increasing concentrations of CDN1163 *vs* the DMSO vehicle (**Fig. 3B**). At high Ca^2+^ conditions, the addition of CDN1163 does not influence the FRET efficiency between SERCA2a and PLN in the presence or absence of AMP-PCP *vs* the DMSO vehicle (**Fig. 3C**). The inability of CDN1163 to compete with PLN is corroborated by a poor fit of the data to a sigmoid curve, i.e., R^2^ values <0.18 when fitted to a non-linear concentration-response regression fit.

**Fig. 3.**
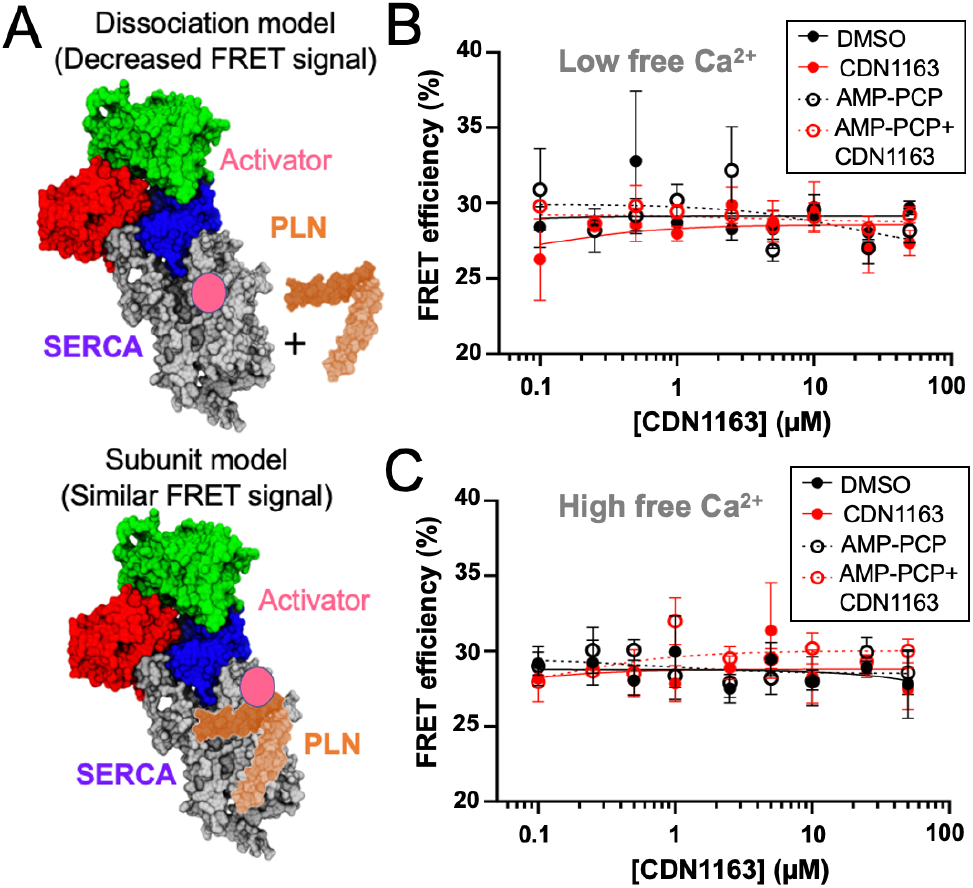
Competitive FRET assays to investigate the effects of CDN1163 on the dissociation of the endogenous SERCA2a–PLN complex. (A) Structural representation of the dissociation and subunit models for activation of SERCA2a by CDN1163 tested in this study. Concentration-response curves at increasing CDN1163 concentrations under (B) low free Ca^2+^ and (C) high free Ca^2+^ conditions. For comparison, we tested the effects of AMP-PCP alone or in the presence of CDN1163. Data are reported as average ± SEM (N=4).

Our primary objective was to assess the concentration-response competition between CD1163 and PLN for binding to SERCA2a. Our analysis did not show a sigmoid relationship between CDN1163 concentrations and the FRET signal between SERCA2a and PLN. However, there may be significant effects on the FRET signal that are independent of the concentration-response fitting model. To test these effects, we performed a two-way ANOVA to establish changes in the FRET signal in response to CDN1163, and the potential effects of Ca^2+^ or ATP on the competition between CDN1163 and PLN compared to the DMSO vehicle (control). The statistical analysis showed that the FRET efficiency between SERCA2a and PLN does not significantly change in the presence of CDN1163 at low (**Fig. 4A**) or high Ca^2+^ (**Fig. 4B**) at all concentrations of the compound, including those near the half activation (i.e., 2.5 µM, **Fig. 4A**) and saturating (i.e., 10 and 50 µM, **Fig. 4A**). Addition of AMP-PCP does not significantly change the SERCA2a– PLN FRET at functionally relevant concentrations of the compound. These findings indicate that CDN1163 does not activate SERCA2a by displacing PLN from the regulatory complex.

**Fig. 4.**
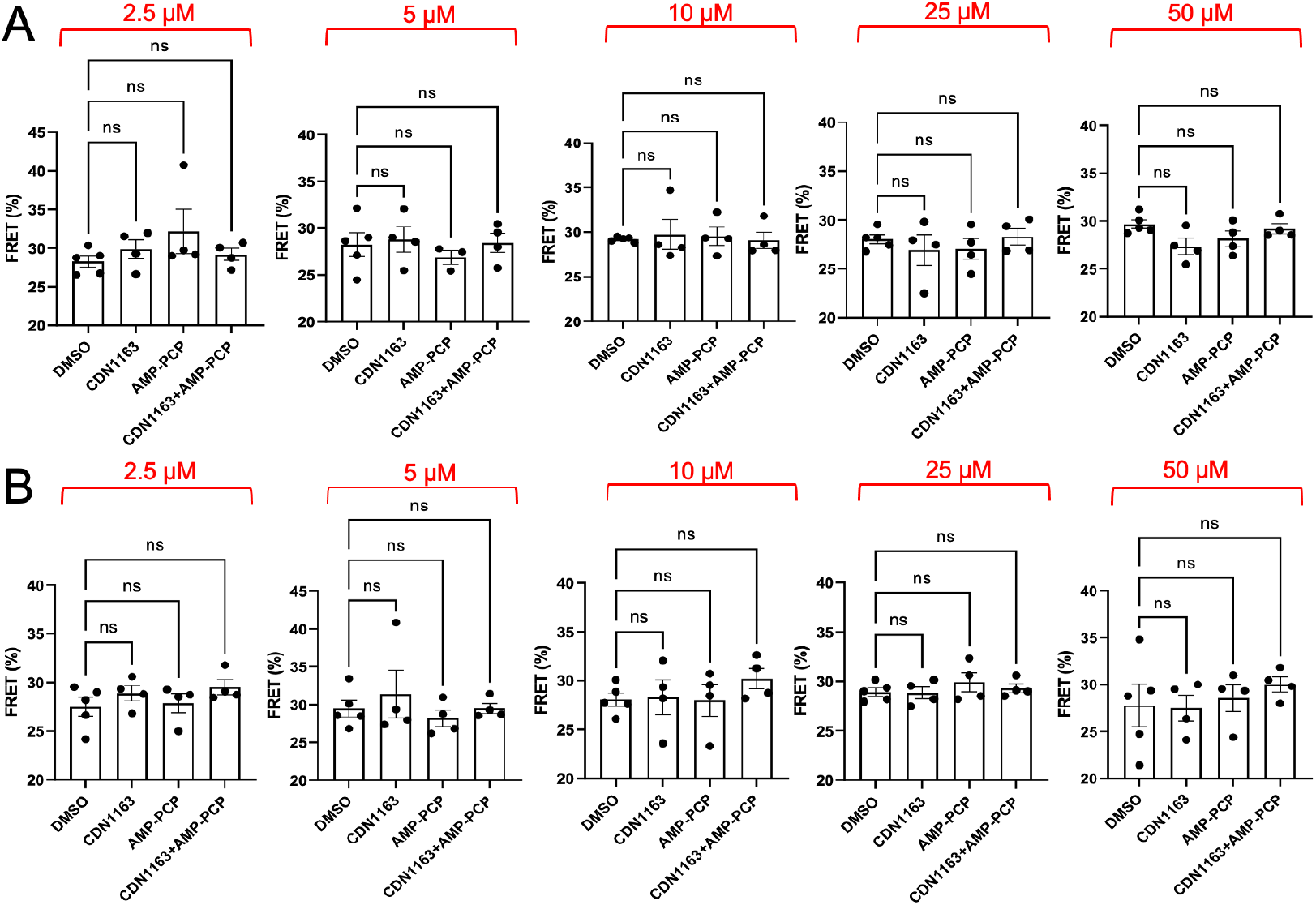
CDN1163 does not induce significant changes in the SERCA2a–PLN interaction. FRET assays performed at (A) low Ca^2+^ conditions (0.01 µM) and (B) high Ca^2+^ conditions (10 µM). In all cases, we found that compared to the DMSO vehicle (control), increasing concentrations of CDN1163 (red numbers) do not significantly change the FRET efficiency between fluorescently labeled SERCA2a and PLN. Addition of the ATP analog AMP-PCP to the SERCA2a-and PLN-containing microsomes does not induce significant changes in the FRET signal vs the control at both low and high Ca^2+^ conditions. Treatment of the microsomes with a fixed concentration of AMP-PCP and increasing concentrations of CDN1163 does not significantly change the FRET efficiency compared to the vehicle, CDN1163 and AMP-PCP treatment groups. Data is shown as mean ± SEM (N=4); we used a two-way ANOVA followed by the Dunnett’s post-hoc test to compare treatments (CDN1163, AMP-PCP, and CDN1163+AMP-PCP) against the DMSO vehicle.

CDN1163 synergizes with AMP-PCP to induce a functional shift in the headpiece populations of SERCA2a

We found that CDN1163 stimulates the ATPase activity of SERCA2a and that dissociation of the endogenous SERCA2a–PLN complex is not the mechanism underlying SERCA2a activation. Previous studies have shown that regulation and Ca^2+^-mediated activation of SERCA2a is achieved through structural shifts of the cytosolic headpiece.^34, 39, 49-52^ Therefore, we tested the hypothesis that CDN1163 allosterically stimulates SERCA2a activity by inducing structural shifts in the cytosolic headpiece of the pump. We note that our combined TCSPC imaging and global analysis approach resolve two primary structural populations of SERCA2a: a catalytically inactive open state, and a closed state that brings together the structural elements required for the formation of a compact headpiece. Indeed, previous studies have shown that the active, disinhibited transported increasingly samples more compact, well-ordered conformations that are catalytically competent.^32, 52-53^ Since our focus is on the structural shifts that correlate with the formation of catalytically competent populations, we therefore represent the structural shifts as the % of the closed population of the headpiece.

In the absence of CDN1163 and at low Ca^2+^ (i.e., [Ca^2+^] <0.1 µM), ∼28% of the SERCA2a populations are in the closed structure. Increasing Ca^2+^ concentrations shift the population of the closed structure of SERCA2a to ∼37% **Fig. 5**); these structural shifts agree with previous studies showing that Ca^2+^ alone is sufficient to redistribute the pump’s structural populations during Ca^2+^ signaling.^34^ Incubation of SERCA2a with either CDN1163 or AMP-PCP induces a shift in the population of the headpiece toward a closed structure, although the effect is more pronounced in the presence of AMP-PCP (**Fig. 5**). Interestingly, the addition of CDN1163 to the SERCA2a/AMPPCP further shifts the population of the headpiece toward a closed structural state that is comparable to that of SERCA2a alone at saturating Ca^2+^ conditions. The cooperative effects of CDN1163 and AMP-PCP are more prominent at Ca^2+^ concentrations between 0.1 and 2 µM. This suggests that CDN1163 synergizes with Ca^2+^ and ATP to induce a structural shift in the cytosolic headpiece of SERCA2a. We tested this hypothesis by performing a two-way ANOVA and the post-hoc Dunnett’s test to compare the changes in the % of the closed headpiece SERCA2a in response to CDN1163, AMP-PCP, and CDN1163/AMP-PCP at twelve Ca^2+^ concentrations.

**Fig. 5.**
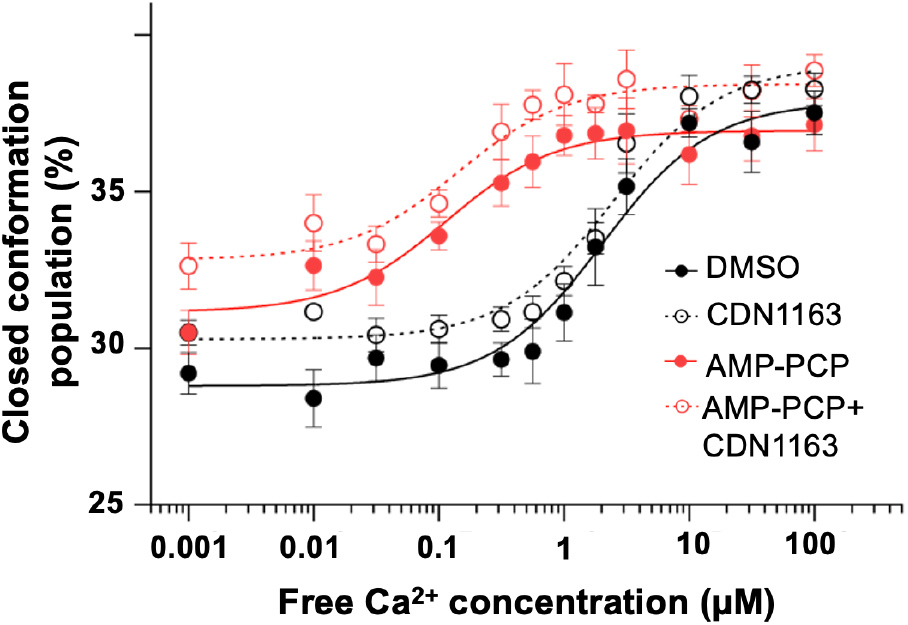
SERCA2a structural change in response to CDN1163. 12-point Ca^2+^ concentration-response curves obtained in the presence of DMSO vehicle (control; solid black trace), CDN1163 (dashed black trace), AMP-PCP (solid red trace) and CDN1163+AMP-PCP (dashed red trace). The structural change in response to ligands is shown as the % of closed population of the headpiece. Data are reported as average ± SEM (N=4).

During cardiac contraction, the intracellular free Ca^2+^ concentration in cardiac cells increases to 1–2.65 µM, allowing interaction between the contractile elements.^54-56^ Relaxation occurs following a decrease in free Ca^2+^ concentrations to 0.1–0.16 µM, causing dissociation of the contractile elements.^54-56^ Therefore, these values represent the upper and lower limits of the range of physiological Ca^2+^ concentrations at which SERCA2a is operational in cardiac cells. The statistical analysis showed that compared to the DMSO vehicle, CDN1163 alone does not have a significant effect on the headpiece population at most Ca^2+^ concentrations tested here (**Fig. 6**). CDN1163 alone significantly shifts the headpiece populations toward a closed conformation (*p*<0.05, **Fig. 6**) at a Ca^2+^ concentration outside the physiological window ([Ca^2+^]=0.001 µM). Compared to the DMSO vehicle, AMP-PCP induces a significant shift in SERCA2a’s headpiece toward a closed population at Ca^2+^ concentrations 0.001, 0.01, 0.03, 0.1 0.56, 1, and 1.8 µM (**Fig. 6**). The effects of AMP-PCP on SERCA2a agree with previous studies showing that the binding of ATP induces closure of the pump’s headpiece.^52^ This effect is not significant at a [Ca^2+^]=0.32 µM, which falls within the physiological window of Ca^2+^ concentrations (**Fig. 6**). AMP-PCP also does not have a significant effect on the head-piece populations at Ca^2+^ concentrations at high Ca^2+^ conditions outside the physiological window (**Fig. 6**).

**Fig. 6.**
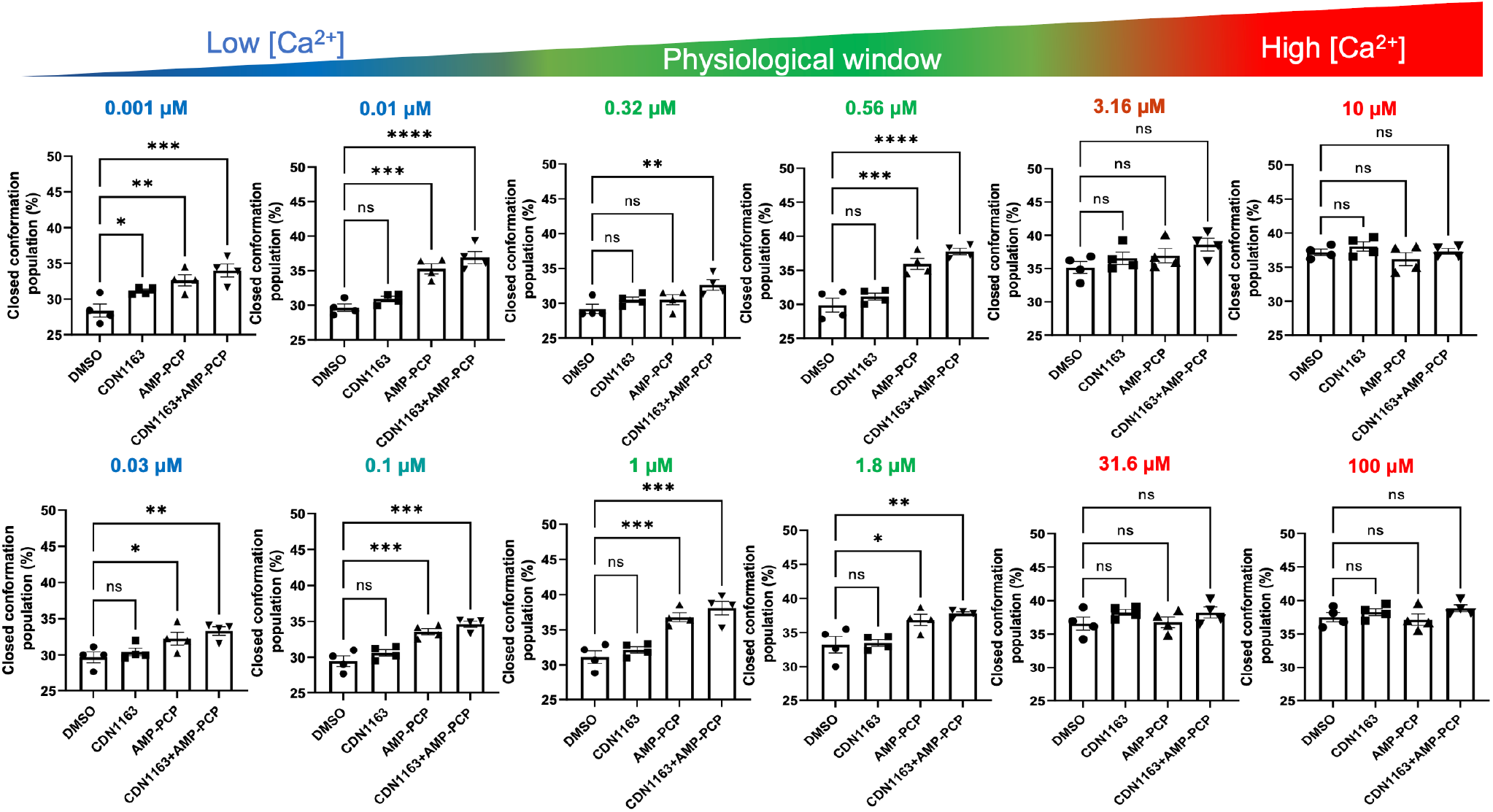
Effects of CDN1163, AMP-PCP, and CDN1163/AMP-PCP on the structural populations of SERCA2a’s headpiece. TCSPC assays were performed at increasing Ca^2+^ conditions from 0.001 µM to 100 µM. Intracellular free Ca^2+^ concentration in cardiac cells fluctuate in the range of 1–2.65 µM and 0.1–0.16 µM during contraction and relaxation of the heart. Therefore, these values represent the upper and lower limits of the range of physiological Ca^2+^ concentrations at which SERCA2a operates. The full range of Ca^2+^ concentrations used in this study is shown, where the physiological window is shown in green, and low and high Ca^2+^ concentrations falling outside this window are shown in blue and red, respectively. In all cases, we used AMP-PCP and CDN1163 concentrations of 0.5 and 10 µM, respectively. Data is shown as mean ± SEM (N=4); we used a two-way ANOVA followed by the Dunnett’s post-hoc test to compare treatments (CDN1163, AMP-PCP, and CDN1163/AMP-PCP) against the DMSO vehicle. \**p*<0.05; \*\**p*<0.01; \*\*\**p*<0.001;**** *p*<0.0001.

The most significant effect on the population shift occurs in the presence of both CDN1163 and AMP-PCP at most Ca^2+^ concentrations below 2 µM, e.g., *p*<0.05 with AMP-PCP *vs p*<0.01 with AMP-PCP+CDN1163 at a free Ca^2+^ concentration of 1.8 µM (**Fig. 6**). The treatment of microsomes with both AMP-PCP and CDN1163 does not have a significant effect on the headpiece populations at high Ca^2+^ concentrations that fall outside the physiological window (**Fig. 6**). Our analysis also showed that CDN1163 and AMP-PCP combined induce a structural shift of the headpiece population that is comparable to that at saturating Ca^2+^ conditions. The combined effect of CDN1163 and AMP-PCP occurs independently of Ca^2+^. For example, we observed similar effects of AMP-PCP *vs* AMP-PCP+CDN1163 treatment groups at Ca^2+^ concentrations 0.01 µM, which falls outside the physiological window, and 0.56 µM, which falls within the physiological window. The ∼10% population shift is comparable to that induced by saturating Ca^2+^, so CDN1163 synergizes with AMP-PCP to populate a closed structure of the head-piece SERCA2a at non-saturating Ca^2+^ concentrations.

### Effects of CDN1163 on the affinity of SERCA2a for Ca^2+^ ions

We showed that CDN1163 potentiates the effects of AMP-PCP to produce a population shift required for the activation of the pump. Besides these structural changes, CDN1163 may activate SERCA2a by increasing its affinity for Ca^2+^. We tested this mechanism first by analyzing the [Ca^2+^]-dependent ATPase activity curves of SERCA2a at compound concentrations between 0.1–100 µM. We found that at concentrations of CDN1163 equal to or less than 1 µM, the compound does not affect the Ca^2+^ affinity for SERCA, illustrating the fact that concentration-response curves do not shift along the x-axis compared to the control (**Fig. 7A**). However, at concentrations of CDN1163 equal or higher than 2.5 µM, the concentration-response curves shift to the right along the x-axis.

**Fig. 7.**
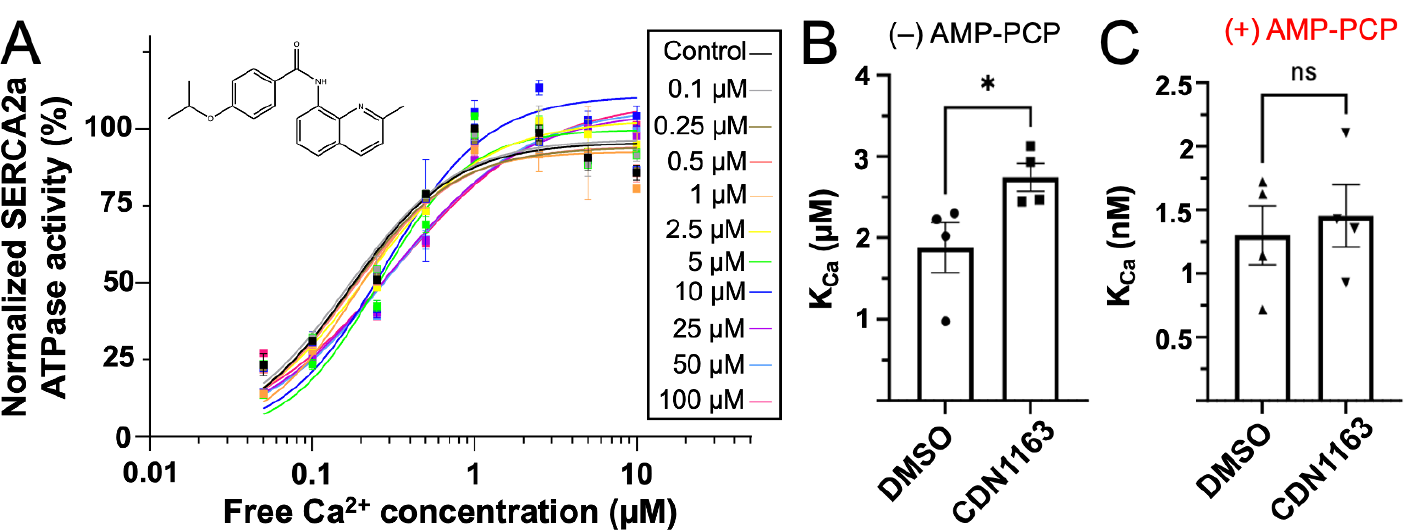
The effects of CDN1163 on SERCA2a’s affinity for Ca^2+^ ions measured by ATPase activity assays and TCSPC imaging. (A) 8-point Ca^2+^ concentration-response of SERCA2a ATPase activity curves obtained at ten concentrations of CDN1163 (0.1 to 100 µM). The activity at each concentration of CDN1163 was normalized relative to the control at [Ca^2+^] =1 µM because this is the point representing the maximal activity of the pump in our assays. Data are reported as average ± SEM (N=3). To eliminate the confounding effects of ATPase hydrolysis, we used TCSPC imaging to measure the effects of 10 µM CDN1163 on SERCA2a’s K*Ca* in the (B) absence and (C) presence of the ATP analog AMP-PCP. Data is shown as mean ± SEM (N=4), and each data point represents a biological replicate. Significance compared to the DMSO vehicle (control) was determined using a two-tailed Mann–Whitney U test; \**p*<0.05.

The ATPase activity curves suggest that while CDN1163 increases the maximal activity of SERCA2a (i.e., turnover rate), it also decreases, albeit modestly, the affinity of the pump for Ca^2+^ (**Fig. 7A**). A limitation of the ATPase activation assays is that the signal depends on hydrolysis of ATP, and therefore, it is impossible to separate the effects of the activator alone or in combination with nucleotide substrate on SERCA2a’s affinity for Ca^2+^. We overcome this limitation by calculating SERCA2a’s affinity for Ca^2+^ directly from the TCSPC imaging curves (**Fig. 5**). The advantage of this approach is that we eliminate the confounding effects of ATPase hydrolysis. We determined the effect of CDN1163 on SERCA’s K_Ca_ at a concentration of 10 µM because it exerts the maximal stimulatory effect in the ATPase activity assays (**Fig. 7A**). The calculated K_Ca_ of SERCA2a in the DSMO vehicle and the absence of AMP-PCP is 1.9 ± 0.3 µM, and treatment of the microsomes with CDN1163 significantly increases the K_Ca_ of the pump to a value of 2.7 ± 0.2 µM (*p*=0.0286, **Fig. 7B**). In the presence of AMP-PCP, the K_Ca_ of SERCA2a is 1.3 ± 0.2 nM and addition of 10 µM CDN1163 decreases the pump’s affinity for Ca^2+^, with a mean K_Ca_ value 1.5 ± 0.2 nM (**Fig. 7C**). These findings agree with the ATPase activity assays and indicate that CDN1163 decreases the SERCA2a’s affinity for Ca^2+^. However, SERCA2a activation requires the presence of nucleotide, and the effect of CDN1163 on K_Ca_ is not significant in the presence of AMP-PCP (*p*=0.8857, **Fig. 7C**). Therefore, we can conclude that CDN1163 does not modulate the affinity of SERCA2a for Ca^2+^ ions at functionally relevant conditions of the pump.

## Discussion

We developed a TCSPC imaging approach to determine the mechanisms for the activation of SERCA2a. This approach, enabled by the recent development of improved fluorescence proteins and hybrid detectors for TCSPC, allows us to detect the individual and combined effects of Ca^2+^, nucleotide substrate, and small-molecule modulators on the structural dynamics of SERCA2a. An advantage of this approach is that we can efficiently perform concentration-response assays to systematically probe the effects of ligands, substrate, and small-molecule effectors on SERCA2a dynamics. An important innovation in this approach is the use of component analysis of globally fitted TCSPC curves to resolve structural transitions within SERCA2a. The powerful combination of TCSPC imaging and component analysis yielded structural insights into mechanisms for SERCA2a activation by a small-molecule modulator in unprecedented detail.

We first used ATPase activation assays to screen for SERCA2a effectors that activate SERCA in a concentration-dependent manner. This is important because there are small-molecule probes that exhibit a “bell” or “U-shaped” behavior, which is indicative of compound-mediated assay interference,^47^ and therefore, renders these molecules unsuitable as molecular probes. Among the SERCA effectors reported in the literature, only CDN1163 acts as a direct activator of SERCA2a, increasing its activity sigmoidally, and with an effect on V_max_ retained at higher concentrations of the compound. These findings agree with a solid-supported membrane biosensing approach showing that CDN1163 enhances SERCA-mediated Ca^2+^ translocation at compound concentrations that are similar to those in this study.^43^ An important question we ask is whether the ∼12% activation of SERCA2a ATPase activity is functionally significant at the cellular level. We have previously used optical mapping experiments to show that CDN1163 significantly increases Ca^2+^ dynamics using human iPSC-derived cardiomyocytes.^48^ These functional effects are comparable to those induced by the adrenergic agonist isoproterenol, which promotes SERCA2a activation via PLN phosphorylation.^14, 57-58^ These studies validate CDN1163 as a probe to systematically characterize the mechanisms for small-molecule activation of SERCA2a.

PLN inhibits SERCA2a by binding to a large pocket in the transmembrane domain of the pump.^59-64^ This interaction populates a Ca^2+^ ion-free intermediate that serves as a kinetic trap that decreases SERCA’s apparent affinity for Ca^2+^ and depresses the structural transitions necessary for Ca^2+^-dependent activation of the pump.^65-67^ Therefore, a conventional concept in the field is that dissociation of the SERCA2a–PLN complex is a requirement for SERCA2a reactivation.^24^ Therefore, we tested whether CDN1163 stimulates the ATPase activity of SERCA2a by displacing PLN from the endogenous SERCA2a–PLN complex. We found that CDN1163 does not dissociate the complex either at functionally relevant or saturating concentrations of the compound. PLN is thought to interact with Ca^2+^-free forms of SERCA and partially dissociate from the pump at high Ca^2+^ concentrations.^68-69^ However, CDN1163 has no effect on the FRET signal between SERCA2a and PLN at saturating Ca^2+^ conditions (i.e., 10 µM). Previous studies have also suggested that PLN and ATP stabilize a functional SERCA–PLN-ATP state that protects the pump against the binding of inhibitors.^70^ We found that in the presence of the ATP analog AMP-PCP at either low or high Ca^2+^ conditions, CDN1163 has no effect on the FRET signal between SERCA2a and PLN, indicating that CDN1163 does not compete with PLN and has no effect on the PLN/ATP-bound state of the pump. This evidence indicates that CDN1163 does not dissociate the SERCA2a–PLN complex, challenging the current paradigm proposing that endogenous PLN needs to be displaced from SERCA to activate the pump.^24-25^

We take advantage of the robustness of the TCSPC imaging method, which has over 4 orders of magnitude difference between signal and noise. We can also perform a global analysis of lifetimes for over 200 fluorescence decays and attribute specific lifetimes to structural states of SERCA2a. The combination of TCSPC and global analysis allows us to resolve the structural states of SERCA2a and determine how these states are redistributed in response to ligands. An important advantage of this approach is that we examine the effects of CDN1163 alone or in combination with Ca^2+^ and nucleotide. We used this approach to demonstrate that CDN1163 does not modulate the affinity of SERCA2a for Ca^2+^ ions at functionally relevant conditions of the pump. Instead, CDN1163 activates SERCA2a by a mechanism in which the activator synergizes with ATP to allosterically redistribute the headpiece populations toward a closed, preactivated conformation of SERCA2a. This population shift occurs only within or below the physiological Ca^2+^ concentrations at which SERCA2a operates.^54-56^ Our combined approach, therefore, suggests a mechanism in which biochemical and structural changes are coupled to allosterically activate SERCA2a.

Crystallography, spectroscopy, and computational studies have shown that activation and inhibition of SERCA correlate with the shifts between open and closed structural populations of the cytosolic headpiece, the domain of the pump that contains the catalytic elements of the pump.^34^ Specifically, activation of the pump occurs through a population shift toward a closed structure of the headpiece, whereas inhibition by small molecules (e.g., thapsigargin) and endogenous PLN occurs through a shift in the headpiece populations toward an open state.^34, 39, 53^ While our study correlates well with this general mechanism, it shows for the first time how these structural changes occur in response to ligands, nucleotide substrate, and an allosteric modulator quantitatively. This is important because TCSPC experiments at all conditions show that SERCA2a’s headpiece undergoes a maximal ∼10% shift in the equilibrium toward the closed state in activating conditions. Relatively small shifts in structural populations are a common theme in the allosteric modulation of sarcomeric proteins. For example, allosteric activation of the smooth muscle myosin induced by phosphorylation of its regulatory light chain occurs with a ∼20% shift in the structural populations within its phosphorylation domain.^71^

Fluorescence spectroscopy studies have suggested that, in the absence of allosteric modulators, the headpiece closure is coupled to substrate ATP binding to the pump.^52^ Our findings agree with these studies and show that AMPPCP induces a significant shift in the headpiece’s population toward a closed conformation at Ca^2+^ concentrations <2 µM. Interestingly, further increases in Ca^2+^ concentration do not result in an additional shift of the headpiece conformation. Instead, the concentration-response curve reaches a plateau at [Ca^2+^]>2 µM independently of AMPPCP. These findings suggest ATP is both a substrate and an effector of SERCA2a. Future studies must be designed to precisely determine the functional effects of Ca^2+^ and nucleotide on SERCA2a.

In conclusion, in this work we introduce TCSPC imaging and global analysis of fluorescent lifetimes, a powerful approach that makes possible the direct analysis of the structural mechanisms underlying allosteric modulation of proteins. A key feature of this combined approach is the resolution of protein structural states and population shifts in response to ligands, substrates, and small molecules. This technical advantage allowed us to monitor the structural mechanisms for activation of the cardiac calcium pump SERCA2a by CDN1163, a validated smallmolecule allosteric modulator. The significance of this space-time resolution is threefold. First, the experiments directly showed that CDN1163 does not compete for binding with SERCA2a’s endogenous regulator PLN and that this effect is independent of Ca^2+^ and ATP. Second, the method allowed us to directly measure the response of SERCA2a to ligands and show that CDN1163 does not activate SERCA2a by modulating the pump’s affinity for Ca^2+^. Finally, we directly resolved structural shifts of SERCA2a to show that CDN1163 synergizes with ATP to redistribute the headpiece populations toward a pre-activated conformation of SERCA2a at non-saturating Ca^2+^ concentrations. This study demonstrates the power of the TCSPC imaging approach, revealing how structural and biochemical changes, defined by bound ligands, are coupled to allosterically activate a therapeutic target in the heart.

## AUTHOR INFORMATION

## Acknowledgments

We thank Gail Rising, Coordinator of the Animal Surgery Operating Rooms at the University of Michigan Unit Laboratory Animal Medicine, for the donation of pig hearts. We thank Sean Cleary for the optimization of Ca^2+^-containing buffers, and Ellen Cho for technical assistance. J.S. was supported by an American Heart Association Postdoctoral Fellowship 830562. This work was supported by grants from the National Institutes of Health R01GM120142 and R01HL148068 (to L.M.E.-F.), and R01HL092321 and R01HL143816 (to S.L.R.). This research was supported in part through computational resources and services provided by Advanced Research Computing, a division of Information and Technology Services at the University of Michigan, Ann Arbor.

## Notes

The authors declare that they have no known competing financial interests or personal relationships that could have appeared to influence the work reported in this paper.

## Data availability

All data are available in the main text.

